# Juvenile hormone degradation enzymes have shared and unique requirements in *Drosophila* development

**DOI:** 10.1101/2025.06.09.657647

**Authors:** Rebecca Spokony, Krystal Goyins, Harry Siegel, Creehan Healy, Isabella Alvarado, Amina Jumamyradova, Yashfa Naseem, Jie Ying, Alexey A. Soshnev, Lacy J Barton

## Abstract

Precise control of hormones is essential for animal development. Hormone bioavailability is regulated by their synthesis, transport, sequestration, natural turnover, and programmed degradation. Here, we use *Drosophila melanogaster* to investigate developmentally programmed degradation of the retinoid-like juvenile hormones (JHs). JHs promote juvenile growth and timely development by functionally opposing the steroid hormone, ecdysone. Despite well-characterized biochemistry, the developmental requirements for programmed JH degradation are poorly understood due to paralogue expansion. To close this knowledge gap, we generated double and triple knockout animals lacking one of two classes of JH degradation enzymes: JH esterases (JHEs) and JH epoxide hydrolases (JHEHs). We found that while both JHEs and JHEHs restrain JH signaling during *Drosophila* juvenile development, each has separate requirements in the regulation of developmental timing and growth. Strikingly, loss of all three JHEHs doubled the length of juvenile development, and the resulting pupae were smaller. Through targeted and genome-wide transcriptome analysis, measurements of hormone-producing glands, and rescue experiments, we uncovered both shared and unique dysregulated gene networks, some of which have established roles in the regulation of both body size and developmental timing. Our comparative analysis of both JH degradation pathways demonstrated that loss of JHEH, but not JHE, activates multiple levels of compensatory feedback to maintain homeostasis within JH and ecdysone axes. These data not only suggest that JHEH-mediated degradation is the dominant programmed JH degradation pathway in *D. melanogaster* development, but they also revealed new JH homeostatic mechanisms more generally. Together, this study provides new genetic tools and insights into programmed hormone degradation.

**Article summary:** Hormones control the timing of developmental transitions and growth. As such, multiple mechanisms ensure precise amounts of a hormone are available at the right time and place. Here, we investigated the poorly understood process of developmentally programmed hormone degradation using juvenile hormones (JHs) in *Drosophila melanogaster* as a model. We generated double and triple knockout strains to disrupt JH degradation and found an unexpected separation of requirements in regulating developmental growth and timing. Through phenotypic and transcriptomic analysis, this study provides insights into the mechanisms and complexities of programmed hormone degradation in animal development.

## Introduction

Development, physiology and reproduction in animals require precise control of hormone signaling. Hormone signaling can be controlled by the presence or absence of receptors, as well as hormone synthesis, transport, sequestration, turnover, and active degradation. How hormones are precisely controlled in space and time remains a significant biological challenge, especially for lipid hormones which are not directly encoded by genes.

Spatiotemporal availability of lipid hormones is largely driven by the balance between synthesis and degradation. A wealth of insights into lipid hormone synthesis has been gained using the model, *Drosophila melanogaster*. Like other insects, *Drosophila* development is driven by fluctuations in two lipid hormones: the steroid hormone, ecdysone, and the retinoic acid-like sesquiterpenoid, juvenile hormone (JH). JH titers are high during larval development and decrease to facilitate ecdysone pulses that allow developmental transitions such as metamorphosis (MORROW AND MIRTH 2024; SHINODA *et al*. 2026). Generally, in holometabolous insects, decreased JH signaling shortens the larval period, resulting in premature metamorphosis, smaller pupae and pupal lethality, while increased JH signaling delays metamorphosis and increases body size (ABDOU *et al*. 2011; MIRTH *et al*. 2014). This potent juvenilizing effect underlies the use of JH mimics to control insect populations around the world (NUR ALIAH *et al*. 2021). Together, these observations highlight the importance of balancing the synthesis and degradation of JH in development.

Whereas much is known about ecdysone, large gaps in knowledge persist for JHs. JH synthesis begins with the conversion of acetyl-CoA to farnesyl-PP through the mevalonate pathway and ends with the downstream JH-specific enzymatic cascade that includes epoxidation of farnesoic acid (JIA *et al*. 2024; FUJINAGA *et al*. 2026), and methylation by JH acid methyltransferase (Jhamt) (NIWA *et al*. 2008). JH synthesis primarily occurs in neuroendocrine corpora allata, which are part of the larval ring gland that also contains the prothoracic gland that produces ecdysone (SHINODA *et al*. 2026). In juvenile and adult animals, JHs circulate through the hemolymph via JH binding proteins, enter responding cells, and bind to transcription factors Methoprene-tolerant (Met) or Germ cell expressed (Gce) to induce transcription-dependent responses (Fig 1a) (JINDRA *et al*. 2015).

**Figure 1:**
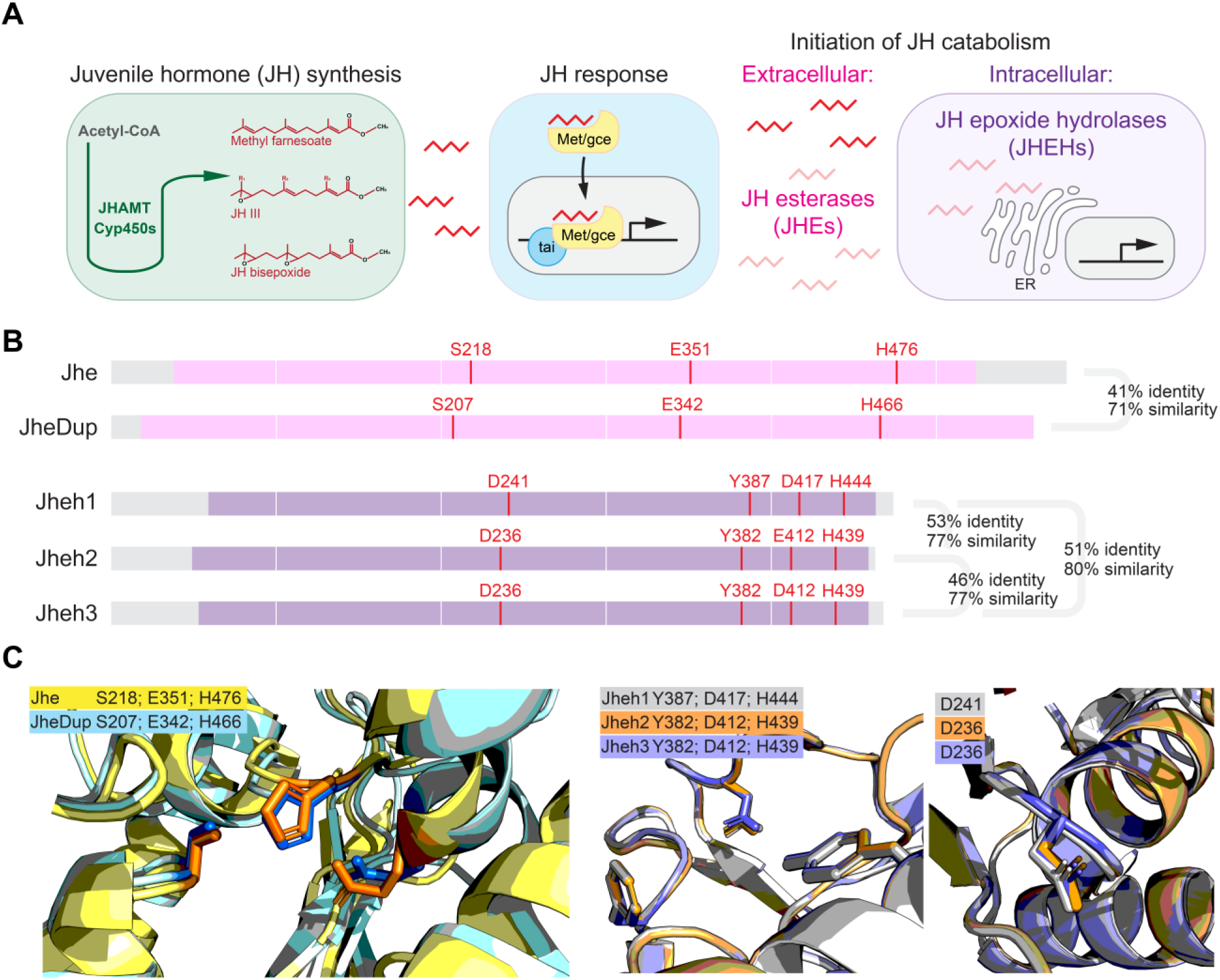
Juvenile hormone degradation enzymes in *Drosophila melanogaster*. **A** Schematic diagram of JH synthesis, response and degradation pathways. *Left*: JH synthesis from Acetyl-CoA precursors through methylation by Jhamt and Cyp450s, Cyp6g2 and Cyp6a13. *Middle*, JH signaling response in which JHs bind transcription factors, Methoprene tolerant (Met) or germ cell expressed (gce), which translocate and bind to response elements together with co-factor, Taiman (Tai). *Right:* JH degradation by intercellular JH esterases (JHEs, pink) to produce JH acids, or intracellular JH epoxide hydrolases (JHEHs, purple) to produce JH diols. **B** Annotated graphic of JHE and JHEH protein features and sequence similarity. Jhe and JheDup share conserved esterase catalytic domains (pink) and Jheh1,2, and 3 share alpha/beta hydrolase catalytic domains (purple). Amino acid involved in catalysis are highlighted in red. **C** Alphafold predicted catalytic pocket of overlaid JH degradation enzyme paralogues with catalytic side chains shown (See Methods for more details).

Juvenile hormones can be passively or actively degraded. Insects’ programmed JH degradation pathways are initiated by two classes of enzymes: JH esterases (JHEs) and JH epoxide hydrolases (JHEHs) (Fig 1a). JHE and JHEH enzymes degrade JHs *in vitro* and reduce JH titers *in vivo* (VENKATESH 1990; CRONE *et al*. 2007; GOODMAN AND CUSSON 2012; C. RIVERA-PÉREZ 2019). Yet, each class of enzyme differs in which JH functional group they modify, as well as when and where they act. JHEs function as monomers and inactivate JHs by removing the methyl group from the carboxylic ester to form JH acid. JHEs generally act extracellularly to modify JHs within the hemolymph circulatory system, though recent work demonstrated that a JHE in *Camponotus floridanus* ants acts intracellularly within the glial blood brain barrier (Fig 1a) (KAMITA AND HAMMOCK 2010; JU *et al*. 2023). The products of JHE action can be used as substrates for Jhamt, thus providing a renewed pool of active JHs (KAMITA AND HAMMOCK 2010). In contrast, JHEHs function as intracellular dimers that irreversibly degrade JHs by hydrolyzing the epoxide groups of JHs or JH acid to generate JH-diol or JH acid-diol (Supp Fig 1, (JINDRA AND NORIEGA 2026)). While the biochemistry of JHEs and JHEHs has been examined *in vitro*, the functional requirements for each JH degradation pathway in development are poorly understood, in part due to paralogue expansion (KONTOGIANNATOS *et al*. 2016; LEBOEUF *et al*. 2018). Indeed, the *D. melanogaster* genome encodes two JHEs, Jhe and Jhe Duplication (JheDup), and three JHEHs, Jheh1, Jheh2, and Jheh3 (CAMPBELL *et al*. 2001; TANIAI *et al*. 2003). Recent work has shown neofunctionalization of *Drosophila* JheDup in olfaction and requirements of Jheh1 and Jheh2 in carbohydrate metabolism (STEINER *et al*. 2017; ROGALSKI *et al*. 2025). Yet, neither JH degradation pathway has been fully disrupted *in vivo* to determine their requirements in development and beyond.

Here, we closed this knowledge gap by generating and characterizing *Jhe, Jhedup* double knockout (DKO) and *Jheh1,2,3* triple knockout (TKO) *D. melanogaster* strains. Using these new genetic tools, we found that loss of each enzyme class differentially affected developmental timing and growth such that JHEs restrain body size, whereas JHEHs are required to ensure timely development in *Drosophila*. Genetic rescue and transcriptomic analyses revealed that JHEs and JHEHs have similar requirements in reducing JH signaling, *cytochrome P450* gene expression, and ring gland size, while JHEHs have unique requirements in the regulation of the ecdysone axis and ribosomal RNAs. Together, our findings provide the first report of the shared and unique requirements for JH degradation enzymes in *Drosophila* development, shedding light on the diversity of mechanisms that control hormone dynamics and homeostasis in development.

## Materials and Methods

### Animal rearing

Stocks used in this study are listed in Supplemental Table 1 and were reared on a molasses/cornmeal medium with methyl 4-hydroxybenzoate (Tegosept, Sigma H3647) as a mold inhibitor. Embryo collections were conducted using cages with removable apple juice plates. Apple juice plates were made of 25% apple juice, 2.5% sucrose, 2.25% Bacto-agar, and 0.15% Tegosept. Experiments were carried out at 25°C, 70% humidity with a 12-hour day-night cycle.

### Predicted Structures

Jhe and JheDup (Uniprot IDs: A1ZA98 and A1ZA97 respectively) predicted structures were retrieved from the AlphaFold Protein Database and created using the AlphaFold Monomer v2.0 pipeline (RRID: SCR_025454). Jheh1,2,3 (Uniprot IDs: Q7JRC3, Q7KB18, Q7K1W4 respectively) predicted dimer structures were retrieved from the SWISS-MODEL repository (RRID: SCR_018123) and were created using ProMod3.

### Generation of double and triple JH degradation gene knock out animals

*JHE DKO* and *JHEH TKO* animals were generated by non-homologous end joining of CRISPR-mediated breakpoints flanking *Jhe* and *Jheh* loci. Guide RNAs were designed based on sequencing relevant PAM sites in the vas-Cas9 injection stock (Bloomington Stock Center 51323). For each knockout, two guides were injected per PAM site for a total of four injected guides each for *Jhe, Jhedup* and *Jheh1,2,3* deletion (Supplemental Table 2). Founders were screened by PCR detection of small products generated by primers flanking predicted cut sites. For the *Jheh1,2,3* locus, six promising founders were identified. For the *Jhe, Jhedup* locus, eight promising founders were identified. For each promising founder, twelve F1 crosses to *Sp, hs-hid/CyO, ft-lacZ* were set up and desired deletions were screened by PCR with genomic DNA templates isolated by QuickExtract DNA Extraction Solution (LCG QE09050) using primers listed in Supplemental Table 2. Precise breakpoints were then identified by Sanger sequencing using the same primers used to screen for deletions.

For sequence-verified *JHEH TKO* lines, 1-6-2; 1-18-8; and 1-19-4, the large deletion encodes a frameshift starting at amino acid 53 of Jheh3, upstream of the hydrolase domain which starts at amino acid 154. Beyond amino acid 53 of Jheh3, peptides of varying lengths (25 for 1-6-2; 93 for 1-18-8; and 193 for 1-19-4) are possible but do not match any *Jheh* sequence. For sequence-verified *JHE DKO* lines 2-1-1; 2-1-3; 2-1-5; 2-1-9; and 2-1-12, the large deletion encodes a frameshift starting at amino acid 62 of JheDup, downstream of the carboxylesterase domain starting at amino acid 30 but upstream of the acetyl esterase/lipase domain at 103 and upstream of the catalytic triad at amino acid 207, 342 and 466. For all lines, there is a potential additional 15 amino acids not matching any Jhe sequence.

### Percent expected class

The percent expected class of zygotic *JHE DKO* and *JHEH TKO* mutants was calculated as done previously (BARTON *et al*. 2014). Briefly, three virgin heterozygous, balanced females were crossed to two heterozygous, balanced males for three days for each of three biological replicates. To allow for sufficient time to eclose, balanced and unbalanced progeny were counted for up to twenty days after the cross was set up.

### Hatch rate

To determine the hatch rate of embryos maternally and zygotically lacking JHEs or JHEHs, less than three-day old homozygous *JHE DKO* and *JHEH TKO* virgin females were crossed to homozygous mutant males in vials for 24 hours alongside control crosses. Then, parents were transferred to egg collection cages for 24 hours before collecting embryos every 24 hours thereafter. Each plate represented one out of three biological replicates. Embryos/eggs were counted upon plate collection and then incubated at 25°C, 70% relative humidity for 36 hours, after which the number of unhatched embryos was counted.

### Developmental timing assays

Parental ages were controlled to mitigate any effects of age on progeny development. Wild type and *JHE(H) KO/CyO, Tubby^1^::RFP* virgin female flies and males were collected for no more than three days before setting up the mating cross. All genetic crosses were set up in vials for 24 hours before transferring parents to embryo collection cages with apple juice plates. Flies were kept in cages for 24 hours to adjust the flies to the collection process after which embryos were collected every six hours. Egg lay was assumed three hours within every six-hour collection window. Because larval density is known to impact pupariation timing in *Drosophila* (KLEPSATEL *et al*. 2018), 75 embryos were placed in each vial containing the same amount of pre-mashed food. Starting on day three after embryo transfer, the number of new Tubby and non-Tubby pupa were quantified every six hours. Pupae were only counted when the prepupal case had formed (P0 or later), and the center of each pupa was marked on the outside of the vial to prevent recounting. Data was collected until five days after the first adult eclosed to avoid quantifying the next generation. Of note, data shown in Figure 3d was obtained during COVID-related laboratory closures with a small incubator which did not hold constant temperature as well as larger incubators used to obtain data shown Figure 6a. While these incubator differences likely account for differences in the severity of developmental delay observed, each experiment is internally controlled with large developmental delays consistently observed.

### Pupal and adult tissue size measurements

Like developmental timing, growth depends upon density of larvae in *Drosophila* (SANTOS *et al*. 1994) (MAGEE AND SPOKONY 2023). To measure pupal length, as well as ring gland sizes, parents were mated in vials for 24 hours, transferred to embryo collection cages for 24 hours and then 25 embryos were placed into each vial. The food in the vials was mashed to allow the larva to feed even in low density conditions. Pharate adult pupae were sexed by the presence of sex combs and imaged using a Nikon SMZ1500 with Nikon Digital Sight 100, a Leica EZ4, or a Leica Investa 3, depending on the experiment.

### RT-qPCR sample collection, RNA isolation, cDNA synthesis, and data analysis

To assess JH and ecdysone signaling across developmental time by qPCR of target genes, wild type, *JHE DKO* and *JHEH TKO* animals were first synchronized by embryonic hatching.

Embryos were collected on apple juice plates and incubated until larvae began to hatch from the eggshell, at which time larvae were cleared from all apple juice plates. Two hours later, newly hatched larvae were transferred to fresh petri dishes containing mushed molasses/cornmeal food until they were 18-, 24-, or 66-hours post hatching, controlling for larval density. Larvae were then phenotyped for the absence of balancers to ensure homozygosity and placed in TRIzol. RNA was extracted using TRIzol (Invitrogen 15596018) followed by isopropanol precipitation. DNA was removed using RQ1 RNase-Free DNase (Promega M6101). cDNA was generated using SuperScript III Reverse Transcriptase (Fisher Scientific 18080-044) using 50% oligo dT and 50% random hexamer primers. qPCR primers used are listed in Supplemental Table 2. qPCR was carried out using PowerUp SYBR Green Master Mix (Thermo Fisher A25742) and a QuantStudio 6 Pro cycler. Fold change was calculated as previously described (TAYLOR *et al*. 2019) and normalized to two reference genes found to have invariant expression across tissues and developmental stages: *Dnm*/*DCTN2-p50 and DCTN5-p25* (CELNIKER *et al*. 2009).

### RNA-sequencing sample collection, RNA isolation, library preparation, and sequencing

To avoid overcrowding, 75 embryos from embryo collection cages were transferred to wide vials. White prepupae were collected from control and mutant genotype vials, which allowed for the pupa to be synchronized to within a 15-minute window of development. Male and female white prepupae were collected and processed separately. Each biological replicate contained 5- 10 larvae. RNA was extracted TRIzol (Invitrogen, ThermoFisher 15596018) followed by isopropanol precipitation. Seven micrograms of total RNA from each replicate were DNaseI-treated and cleaned up using Monarch Total RNA Miniprep Kit (NEB T2010S). One microgram of total RNA was used for RNA-Seq library preparation, using NEBNext Ultra II Directional RNA Library Prep Kit for Illumina (NEB E7760) with NEBNext Poly(A) mRNA Magnetic Isolation Module (NEB E7490). Samples were indexed with unique dual index primer pairs from NEBNext Multiplex kit (NEB E6440) and processed according to the manufacturers’ instructions, with 8 cycles of amplification. Individual libraries were sized and quantified using Tapestation 4200 and High Sensitivity D1000 ScreenTape (Agilent 5067-5584), and Qubit fluorometer with dsDNA HS Assay kit (Thermo Q32854). Equimolar amounts of individual libraries were combined in one pool and sequenced by a commercial provider (Azenta Inc.) using 150 bp paired-end chemistry on NovaSeq X Plus with 5% phiX spike-in.

### RNA sequencing analyses

Demultiplexed .fastq files were aligned to BDGP dm6 using HISAT v 2.2.1 (RRID: SCR_015530 (KIM *et al*. 2019)), converted to .bam, sorted and indexed using SAMtools 1.21 view (RRID: SCR_002105), sort and index commands with default flags (DANECEK *et al*. 2021), and visualized as .bigwig using deepTools 3.5.6 bamCoverage “--binSize 1” flag (RRID: SCR_016366 (RAMIREZ *et al*. 2016)). Read quality and alignment statistics were assessed with FastQC and multiQC (RRID: SCR_014982 (EWELS *et al*. 2016)). Read counts were summarized from sorted .bam files using HTSeq 2.0.3 “htseq-count” command using BDGP dm6.46 113 release .gtf reference (RRID: SCR_011867 (PUTRI *et al*. 2022)). Differentially expressed genes were identified using DESeq 2 (RRID: SCR_015687 (LOVE *et al*. 2014)), with thresholds of fold change >1.5 and p value <0.05 adjusted for multiple testing using Benjamini-Hochberg correction. Raw reads, counts, and .bw tracks are deposited to Gene Expression Omnibus under accession number GSE286242. The functional enrichment analysis was performed using g:Profiler (version e111_eg58_p18_f463989d) with g:SCS multiple testing correction method applying significance threshold of 0.05 (RRID: SCR_006809 (KOLBERG *et al*. 2023)).

### Immunohistochemistry

Wandering third instar larvae were collected from six replicate vials seeded with 75 embryos each as described elsewhere. Male and female ring glands were dissected according to (IMURA *et al*. 2017). Briefly, the anterior third of the larvae were dissected and inverted, then fixed in 0.4% PFA for 30 minutes. Central nervous systems and ring glands were dissected and stained for DAPI (1.0ug/mL, Invitrogen D1306) for 20 minutes in PBS. Images were collected using a Dragonfly 200 spinning disk confocal (Andor/Oxford Instruments) at 20x magnification and two-micron step size. Imaris (RRID: SCR_007370) was used to measure PG and ring gland volume. Two machine learning models were trained on control images: one to identify ring glands and another for prothoracic gland nuclei. Surfaces were generated using these models with manual refinement, and the volume of ring gland and prothoracic gland nuclei were obtained. Adobe Illustrator was used to make all figures (RRID: SCR_010279).

### Ecdysone rescue experiments

Wild type and *JHEH TKO* embryos were collected from four-hour egg lays to ensure synchronous development. Animals were then left on apple juice plates supplemented with ample mashed molasses/cornmeal food for 72 hours. After 72 hours, 50 larvae were transferred to each vial, containing either 0.5mg/g of 20-Hydroxyecdysone (20E, Sigma H5142, dissolved in 100% Ethanol) or the equivalent amount of 100% ethanol. All experiments contained three biological replicates. To measure time-to-pupariation, pupae were quantified every 12 hours until five days after the first adult eclosed to avoid quantifying the next generation. To measure eclosion rates, we separated pupae into 96-well plates and quantified adult animals after 10 days. Pupal length was measured as described elsewhere in these methods.

### Statistics

Normality and variance of samples were assessed using Shapiro and Levine tests in R (RRID:SCR_001905), T tests were used when data was parametric and a Mann–Whitney U test was used when data was non-parametric, as noted in the figure legends. For the developmental timing experiments, the time at which 50% of the larva had pupariated was found using linear interpolation between the nearest time points.

## Results

### JHEs and JHEHs restrain JH signaling during *Drosophila* development

Due to paralogue expansion, neither JH degradation pathway has been fully disrupted in *D. melanogaster*. Indeed, despite 71-80% similarity and 41-53% identity (Fig 1b), the predicted tertiary structures and binding pockets are nearly identical (Fig 1b, c, Fig S1). Since each paralogue has the required catalytic motifs, to fully abolish each JH degradation pathway, we generated deletions in *Jhe*, *Jhedup* and *Jheh1, Jheh2, Jheh3* by CRISPR gene editing (Fig 2a). We screened founders by PCR using primers flanking desired cut sites and recovered five lines lacking *Jhe* and *Jhedup* originating from a single founder, which will be henceforth referred to as *JHE DKO*. We also generated three lines lacking *Jheh1, Jheh2, Jheh3* from three separate founders, which will be referred to as *JHEH TKO* for JHEH triple knockout. Each line was sequenced for precise breakpoints. While all *JHEH TKO* lines retained the *Jheh3* promoter, no edited genomes encode functional Jhe and JheDup, or Jheh1, Jheh2, and Jheh3 (Fig 2a).

**Figure 2:**
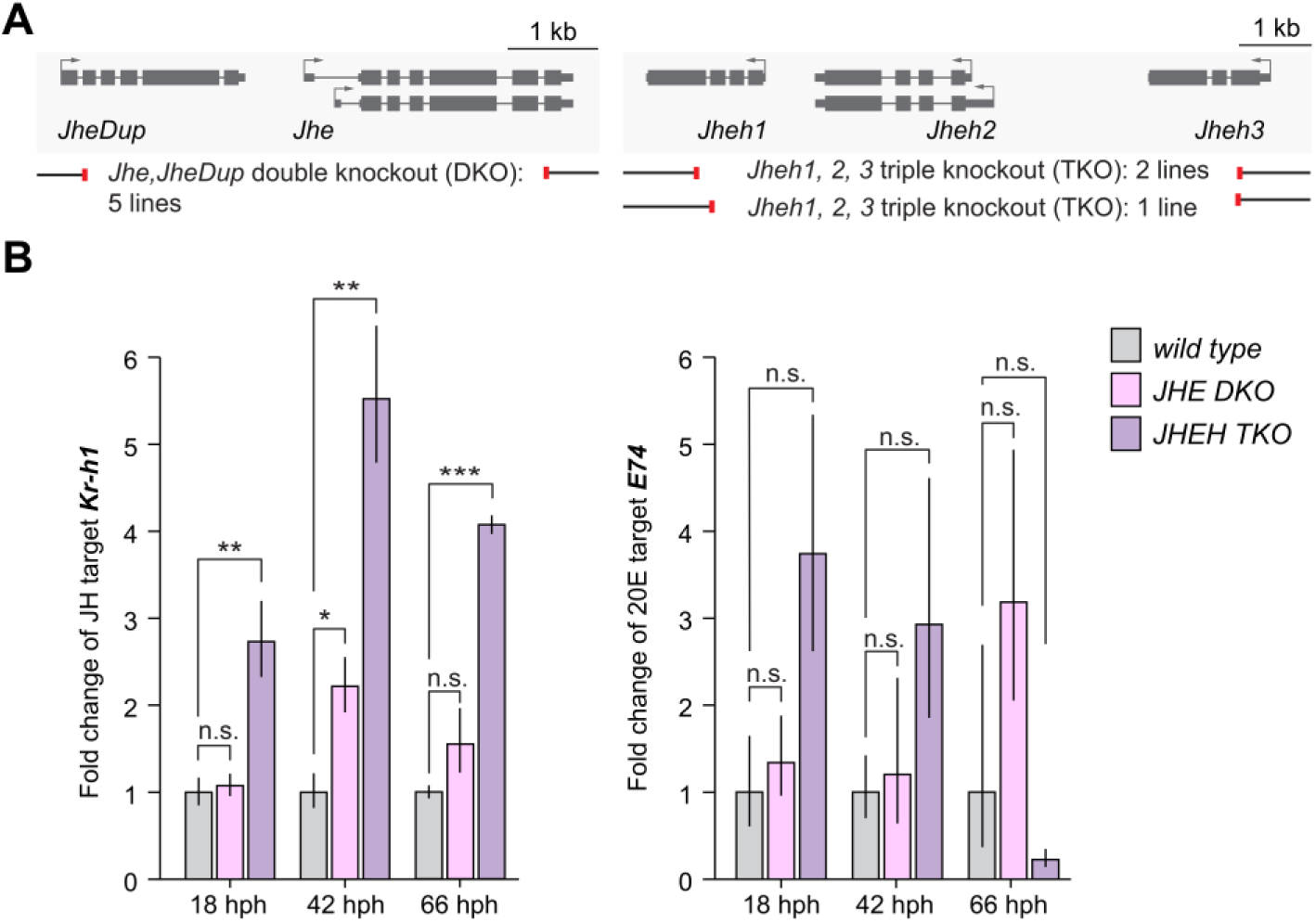
Both JHEs and JHEHs restrain JH signaling in *Drosophila*. **A** Schematic of genes encoding JH degradation enzyme and knockout strains generated in this study. Exons are shown as grey rectangles. Guide RNAs used to delete both *Jhe* and *Jhedup* and all three copies of *Jheh* in respective mutants are shown underneath each locus as red lines. **B** Relative mRNA expression levels of JH target, *Kr-h1* (*Left*), and ecdysone target, *E74 (Right)*, as determined by RT-qPCR of mRNA from whole larvae. Three separate timepoints were examined: 18-, 42-, and 66-hours post-hatching (hph). Data are presented as mean +/- SEM and analyzed for significance using a two-tailed t-test assuming unequal variances (* = p<0.05, ** = p<0.01, *** = p<0.001, n.s. = not significant).

To identify developmental requirements, we first assessed the viability of animals lacking JHEs or JHEHs. To avoid confounding effects of second site mutants, we crossed our newly generated *JHE DKOs* to a deficiency line lacking both *Jhe* and *Jhedup*. At the time of these studies, a deficiency line lacking all three JHEHs was not available. Thus, we crossed two newly generated *JHEH TKO* lines from two independent founders to each other. Under non-crowding laboratory conditions, an expected percentage of heterozygous and homozygous mutant adult progeny eclosed into adults (Fig S2a). To test whether maternally provided JHEs or JHEHs contribute to embryonic development, we generated embryos lacking maternal and zygotic JHE or JHEHs and found that the percentage of embryos lacking either class of JH degradation was comparable to wild type levels (Fig S2b). Together, these data indicate that neither JHEs nor JHEHs are required for viability in *D. melanogaster*.

To assess whether JHE- or JHEH-mediated JH degradation pathways regulate JH or ecdysone signaling, we measured mRNA levels of the classic JH target, *Krüppel homolog 1* (*Kr-h1*) and the ecdysone target, *E74*, by qPCR. To precisely control for developmental time, we transferred newly hatched larvae to vials and then isolated RNA from larvae 18 hrs, 42 hrs and 66 hrs later. We found that *Kr-h1* expression was significantly increased in *JHE DKO* animals at 42 hrs post hatching, which correlates to the second half of second instar, but not 18 or 66 hours post hatching (Fig 2b). In contrast, *Kr-h1* was significantly increased in *JHEH TKO* animals across all timepoints. Interestingly, *E74* mRNA levels were not significantly dysregulated in the absence of either JH degradation pathway (Fig 2b). Together, these data demonstrate that both JHE and JHEH pathways restrain JH signaling during partially overlapping periods of larval development.

### JHEs and JHEHs have opposing requirements in developmental growth and timing

While collecting larvae at precise hours post-hatching to measure JH and ecdysone signaling targets, we noticed that the size of *JHEH TKO* larvae was remarkably heterogenous. This variability could be due to either dysregulated larval growth and/or developmental timing, as elevated JH titers coincide with larger body size and an extended juvenile state in many insects (WHEELER AND NIJHOUT 1981; NIJHOUT *et al*. 2006; MIRTH *et al*. 2014; ZHANG *et al*. 2018). To determine whether either JH degradation pathway regulates growth, we measured pupal length in *JHE DKO*, *JHEH TKO* and wild type animals. As *Drosophila* size is highly sensitive to animal density during development, we transferred 25 embryos to individual vials (SANTOS *et al*. 1994; MAGEE AND SPOKONY 2023). Consistent with the role of JH function in larval growth, JHE DKO animals were larger than wild type controls across both sexes (Fig 3a,b). In contrast, JHEH TKO animals were significantly smaller (Fig 3a,b). To determine whether differences in pupal length were due to loss of JHEs or JHEHs, we generated *JHE DKO* and *JHEH TKO* animals that carried exogenous Bacterial Artificial Chromosomes (BACs) containing either both *Jhe, Jhedup* loci, or all three *Jheh1, 2, 3* genes. While the *JHE DKO* and *JHEH TKO* lines were mutant for the *white* gene, BAC rescue animals were *white+*. Thus, to control known defects associated with *white* loss (GALLONI AND EDGAR 1999), we included a second control, Oregon R crossed to *w^1118^* (OreR x *w^1118^*), for this and all subsequent experiments. Quantification of pupal length revealed that the increased body size was partially rescued in *JHE DKO, BAC[Jhe, Jhedup]* females but not males. In contrast, the smaller body size was fully rescued in *JHEH TKO, BAC[Jheh1,2,3]* animals of both sexes (Fig 3a,b). Together, these data indicate that *Drosophila* JHEs and JHEHs have different requirements in developmental growth.

**Figure 3:**
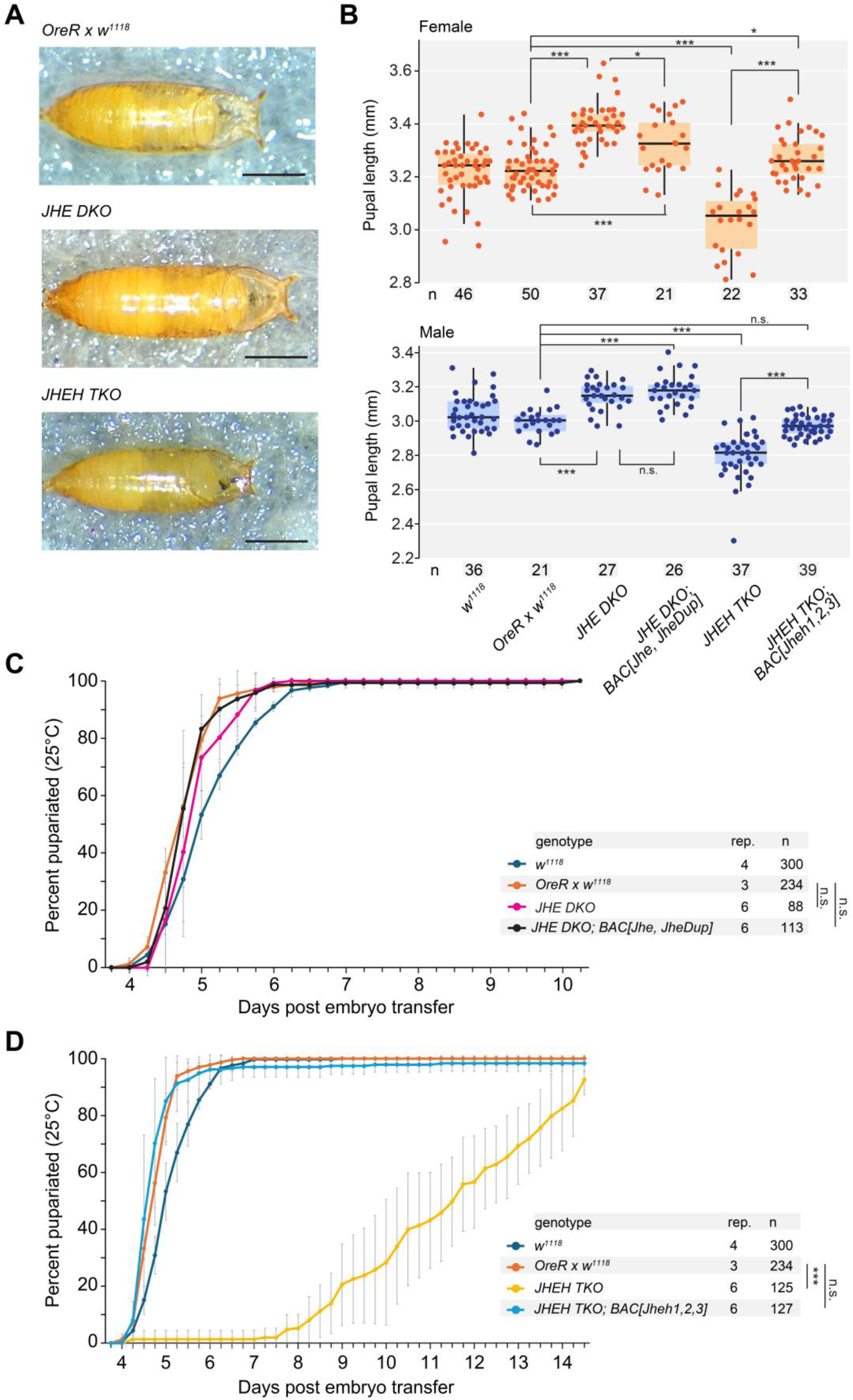
JHEs and JHEHs have opposing requirements in developmental timing and growth. **A** Representative images of wild type, *JHE DKO*, and *JHEH TKO* pupae, scale bars represent 1 mm. **B** Graph of pupal length in females (orange, top) and males (blue, bottom) in each. Genotype is noted under the male graph. Each dot represents one pupa, with total sample size noted below each box plot. Box plots represent median, 25^th^ through 75^th^ percentile with 1.5 IQR whiskers. For B and C, a Mann-Whitney U test was used to test significance (* = p<0.05, ** = p<0.01, *** = p<0.001, n.s. = not significant). **C** Graph of the percent pupariation relative to days post egg lay in control animals, *JHE DKO* animals, or *JHE DKO* animals carrying a BAC that contains both *Jhe* and *Jhedup* loci. **D** Graph of the percent pupariation relative to days post egg lay in control animals, *JHEH TKO* animals, or *JHEH TKO* animals carrying a BAC that contains *Jheh1*, *Jheh2*, and *Jheh3* loci. For C and D, all 50% pupariation data was normally distributed and used for a two-tailed Student’s t-test was used to test significance (* = p<0.05, ** = p<0.01, *** = p<0.001, n.s. = not significant). All experiments were performed at 25°C. Each dot indicates mean percent pupariated and whiskers represent standard deviation. The number of biological replicates (rep) and total number of animals (n) are shown to the right of the genotype.

To determine if either JH degradation pathway is required for proper developmental timing, we quantified the hours it took *JHE DKO*, *JHEH TKO*, and control animals to pupariate. To control the effect of animal density on developmental timing (KLEPSATEL *et al*. 2018), we transferred 75 embryos laid over four hours to individual wide vials and quantified new pupae every six hours until five days after the first adult progeny eclosed to avoid quantifying the F2 pupae. *JHE DKO* animals displayed no delay in pupariation relative to controls (Fig 3c). In contrast, it took seven days longer for *JHEH TKO* animals to pupariate relative to control animals such that 50% of control animals pupariated 4.5 days after embryo transfer while 50% of *JHEH TKO* animals pupariated after 11.5 days (Fig 3d). To determine if any observed prolonged juvenile stages were due to loss of *Jheh* genes, second site mutations, or a gain-of-function phenotype from the intact *Jheh3* promoter (Fig 2a), we measured pupariation timing and found that *JHEH TKO, BAC [Jheh1,2,3]* animals pupariated at the same time as control animals, indicating that the prolonged juvenile stage is due to loss of JHEHs (Fig 3d). Together, these data suggest that JHEHs are required to maintain normal developmental timing in *D. melanogaster*.

### JHE and JHEH degradation enzymes have shared and unique impacts on the transcriptome

Our results thus far indicate that JHEs and JHEHs have separate requirements in the regulation of body size and developmental timing. To understand the nature of these requirements, we profiled transcriptomes of *JHE DKO*, *JHEH TKO* and control males and females by bulk RNA sequencing. Given the dramatic developmental delay in *JHEH TKO* animals, we isolated RNA from animals within the 15 minute-long white prepupae stage to ensure comparison of equivalent developmental stages (BAINBRIDGE AND BOWNES 1981). Principal component analyses of three biological replicates per sex per genotype highlighted a high degree of similarity within *JHE DKO* and *JHEH TKO* biological replicates, both of which clustered separately from wild type samples (Fig S4a). While more variability was observed between sexes than between genotypes, many differentially regulated genes (DEGs) were detected. *JHE DKO* animals had a total number of 1,808 DEGs, while *JHEH TKO* animals had a total of 2,875 DEGs, consistent with their more severe developmental phenotypes (Fig 4a). As expected based on previous studies demonstrating that excess of the JH target, *Kr-h1*, blocks entry into metamorphosis (MINAKUCHI *et al*. 2008; ZHANG *et al*. 2018), we did not detect increased *Kr-h1* in JHE DKO nor JHEH TKO animals by the white prepupal stage. The general direction of altered gene expression was also different, with slightly more genes downregulated compared to upregulated in *JHE DKO* animals of both sexes, while the reverse was observed in *JHEH TKO* animals with more genes upregulated compared to downregulated in both sexes (Fig 4a).

**Figure 4:**
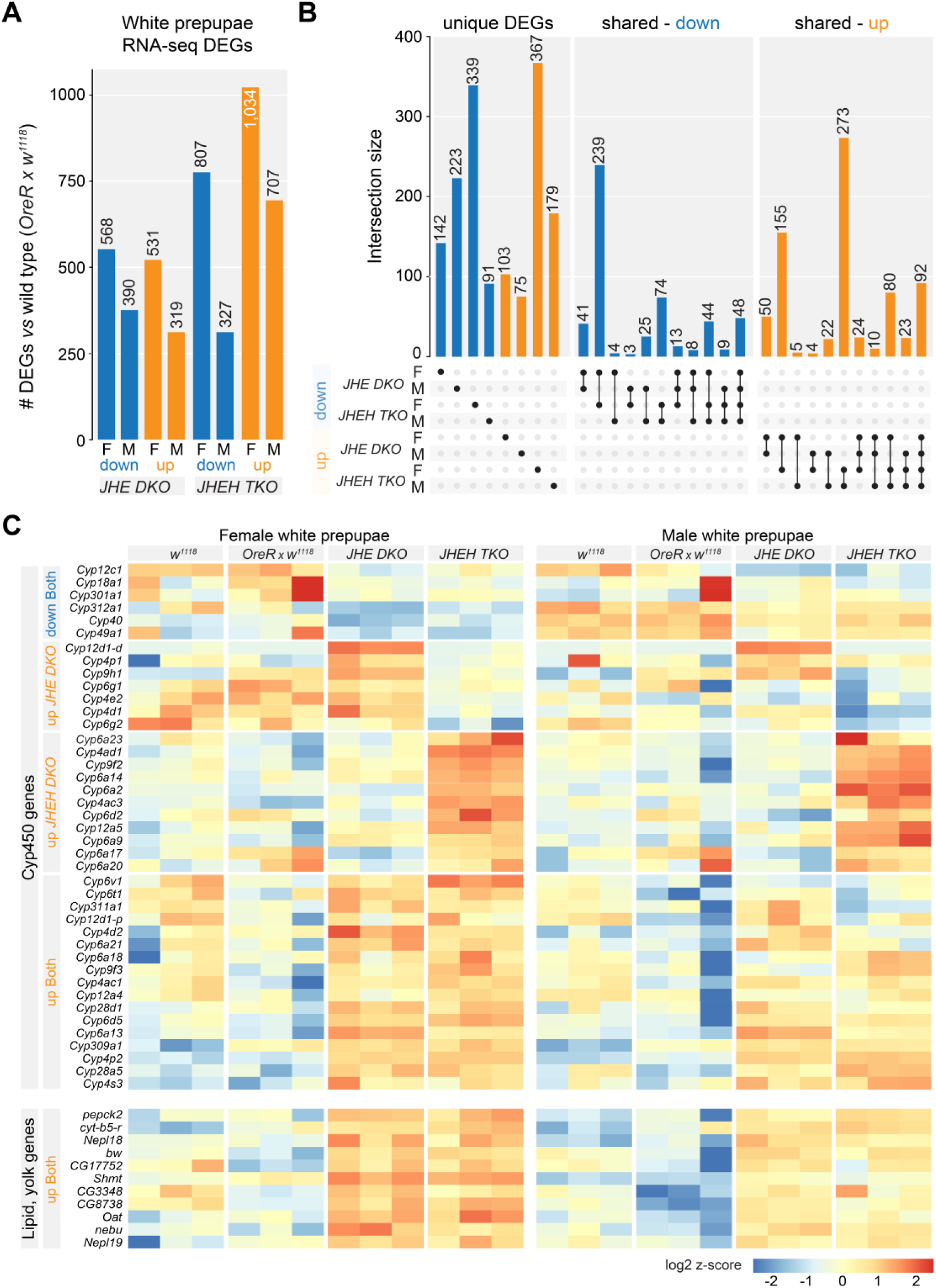
JHEs and JHEHs have shared and unique impacts on the pre-metamorphic transcriptome. **A** Graph of the total number of Differentially Expressed Genes (DEGs) from bulk RNA sequencing of white prepupae. Downregulated genes are in blue and upregulated genes are in orange. The total number of DEGs is shown at the top of each bar. **B** UpSet plot showing either DEGs that are uniquely dysregulated in either *JHE DKO* or *JHEH TKO* animals (*Left*), or DEGs shared by *JHE DKO* and *JHEH TKO* animals (*Right*). The genotypes are noted below, and which samples display the shared DEGs is noted by solid circles with a connecting line. The number of DEGs is shown at the top of each bar. **C** Heatmap of differentially expressed Cytochrome P450 (*top*) and fat body and yolk nuclei genes (*bottom*). Genes are noted to the left, genotype and sex (M or F) is noted above each column, and heat map scale is below.

Analysis of the total number of DEGs revealed more dysregulated genes in both *JHE DKO* and *JHEH TKO* females than males of the same genotype relative to control animals (Fig 4a). Moreover, *JHE DKO* and *JHEH TKO* females shared a substantial number of overlapping downregulated (42%) and upregulated (29%) DEGs, while males had very little shared DEGs (Fig 4b, Datasets S1 and S2). Despite these differences, 14% of all downregulated DEGs and 29% of all upregulated DEGs were shared among males and females of both *JHE DKO* and *JHEH TKO* animals (Fig 4b).

We next performed gene ontology analysis to gain insights into shared and unique requirements of JHE and JHEH degradation pathways (Dataset S3). Consistent with dysregulated JH signaling, there was a significant enrichment of fat body and yolk nuclei genes upregulated in both *JHE DKO* and *JHEH TKO* animals (Fig 4c, (WANG *et al*. 2017)). Also shared upon loss of either JH degradation pathway was dysregulation of genes encoding Cytochrome P450s (Cyp450s) (Fig 4c) and glutathione metabolism, including several Glutathione S-transferase (GST) genes (Supp Fig 4b). These gene products are involved in the production and degradation of hormones, metabolites and xenobiotics and have been found to be induced by exogenous JH (FEYEREISEN 1999; WU AND LU 2008). *JHEH TKO* animals had the greatest amount of Cyp450 dysregulation, with 22 out of the 83 total Cyp450s genes in the *Drosophila* genome and 26 out of 80 total glutathione metabolism genes in the *Drosophila* genome were upregulated (Fig 4c, Supp Fig 4b). Nonetheless, ten Cyp450s and ten glutathione metabolism genes were upregulated upon loss of either JHEs or JHEHs (Fig 4c, Supp Fig 4b). Together, these findings suggest the same JH-linked gene regulatory networks are impacted by loss of either JHEs or JHEHs at the end of the larval period. In addition to these shared DEG signatures, gene ontology also revealed unique DEG signatures. For example, 39 spermatogenesis-related genes were downregulated in *JHE DKO* males (Fig S4c). In contrast, downregulated DEGs in *JHEH TKO* animals of both sexes were significantly enriched for genes involved in iron binding, small molecule transport, metabolism, seminal vesicle biology, and ribosomal biology (Dataset S3 Fig 6c). Together, this transcriptome-wide analysis revealed shared and unique requirements for JHE and JHEH enzymes in *D. melanogaster* development.

### JHEHs are integrated into multiple homeostatic feedback mechanisms

Further analysis of the transcriptomic profiles in *JHE DKO* and *JHEH TKO* animals revealed new signs of homeostatic feedback within JH/ecdysone axes. Several genes encoding ecdysone synthesis genes were upregulated in *JHEH TKO* animals of both sexes (Fig 5a, b). Interestingly, RNA from the intact *Jheh3* promoter was upregulated 13-fold in *JHEH TKO* animals (Fig 5b). Together, these findings suggest that loss of JHEHs activate autoregulatory mechanisms as well as compensatory feedback to increase ecdysone synthesis.

**Figure 5:**
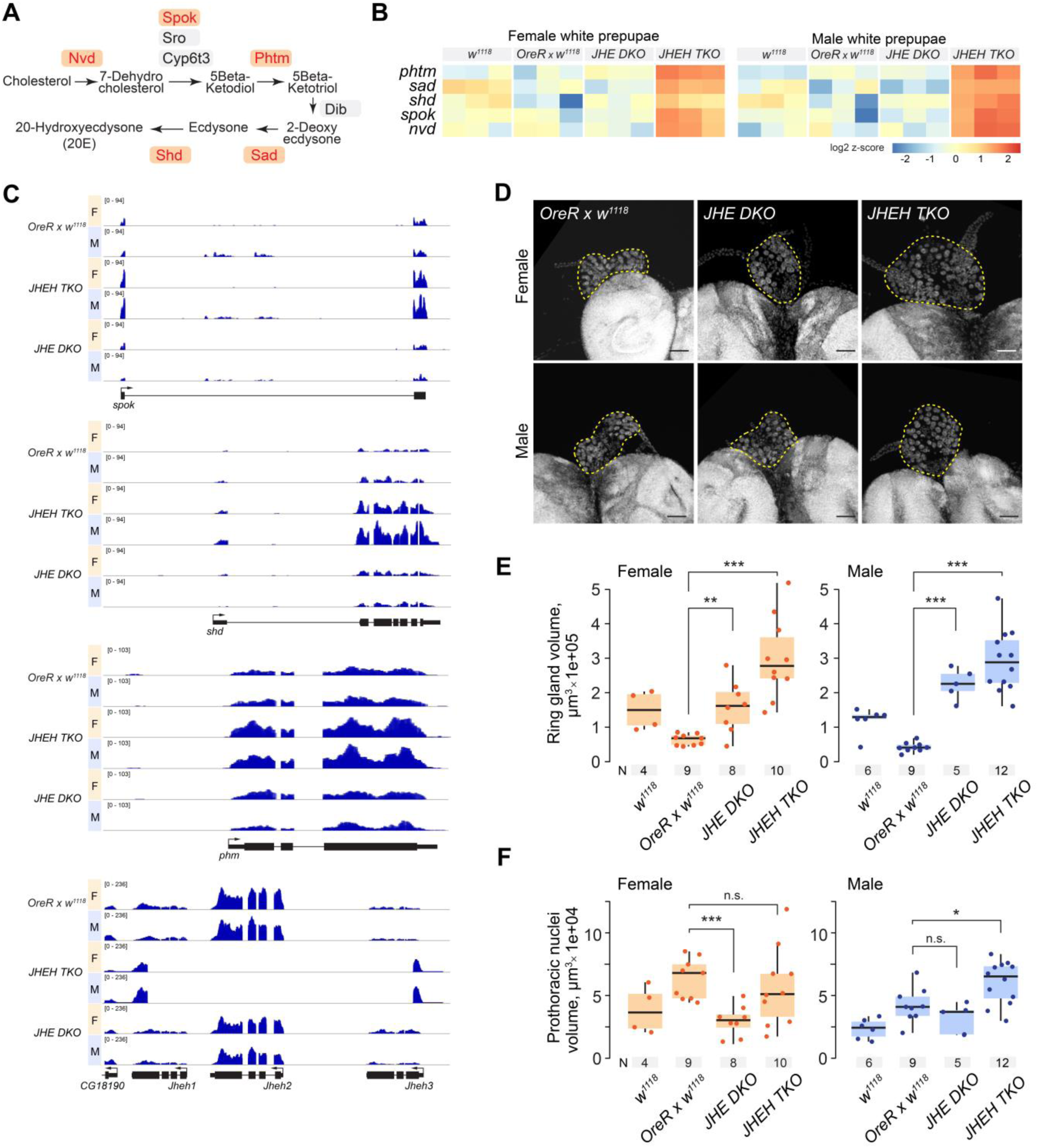
JHEH loss induces compensatory feedback at multiple levels to maintain ecdysone/JH homeostasis. **A** Schematic of ecdysone synthesis cascade. **B** RNA-sequencing tracks showing reads for ecdysone synthesis genes, *spok*, *shd*, and *phm*, as well as *Jheh1*, *Jheh2*, and *Jheh3* loci. Genotype is noted to the left, the gene locus is shown below the tracks. M=male, F=female. **C** Heatmap of differentially expressed ecdysone synthesis genes in white prepupae detected by RNA-sequencing. Genes are noted to the left, genotype and sex (M or F) is noted above each column, and heat map scale is below. **D** Representative images of ring glands from wandering third instar larval females (*top*) and males (bottom) stained with DAPI (gray). Ring glands are outlined with a dashed line. Scale bars represent 50 mm. **E** Quantification of ring gland volume in wandering third instar females (orange, left) and males (blue, right). **F** Quantification of the total volume of prothoracic gland nuclei in each ring gland in wandering third instar females (orange, left) and males (blue, right). For E and F, dots indicate the measurement of an individual ring gland and sample size for each group is shown under the respective box plot. Box plots represent median, 25^th^ through 75^th^ percentile with 1.5 IQR whiskers. A Mann–Whitney U test was used to test significance (* = p<0.05, ** = p<0.01, *** = p<0.001, n.s. = not significant).

Ecdysone is produced by the prothoracic gland, which contributes the majority of the volume to the larval ring gland (KING *et al*. 1966). To determine whether transcriptional signatures of feedback are linked to changes in the ring gland, we dissected them from *JHE DKO*, and *JHEH TKO*, and control wandering third instar larvae and stained them for DNA (Fig 5d).

Quantification of ring gland volume revealed both *JHE DKO* and *JHEH TKO* ring glands were significantly larger than wild type controls despite the smaller body size observed in *JHEH TKO* animals (Fig 3, Fig 5e). Ecdysone synthetic capacity scales with the number of endocycles in the prothoracic gland (OHHARA *et al*. 2017). To estimate endocycling status, we measured total prothoracic gland nuclei volume using DAPI signal and found DAPI volume per prothoracic gland was not consistently increased or decreased across sexes or genotypes (Fig 5f), suggesting the increase in ring gland size in both *JHE DKO* and *JHEH TKO* animals is not due to increased endocycling. Together, these data indicate that increased expression of ecdysone synthesis genes is linked to increased size, but not endocycling, of ecdysone-producing prothoracic glands in *JHEH TKO* animals.

### Defects in developmental timing, but not growth, in the absence of JHEHs is linked to dysregulated ecdysone signaling

Increased or prolonged JH signaling prevents ecdysone peaks that drive metamorphosis (NIJHOUT AND WILLIAMS 1974; ZHANG *et al*. 2018). Our findings demonstrate that while both JHE and JHEH degradation pathways restrain JH titers during at some stages in larval development, loss of JHEH uniquely causes a strong developmental delay that cannot be overcome by the increased expression of ecdysone synthesis genes or increased prothoracic gland size observed in these animals. Together, these data suggest that delayed pupariation in JHEH animals is due to insufficient ecdysone signaling. To test this hypothesis, we fed wild type and *JHEH TKO* larvae the active form of ecdysone, 20E, and measured time to pupariation.

Strikingly, feeding exogenous 20E partially but significantly rescued delayed pupariation in *JHEH TKO* animals (Fig 6a). However, further analysis revealed that while *JHEH TKO* animals fed 20E pupariated faster, rescue of developmental timing led to smaller animals than *JHEH TKO* animals fed control ethanol (Fig 6b). Moreover, almost half of 20E-fed *JHEH TKO* animals failed to eclose into adults, a phenomenon that did not occur in 20E-fed wild type animals (Fig S6). Together, these data suggest while delayed metamorphosis in the absence of JHEH is due to insufficient ecdysone, JHEH likely has additional functions in growth. Indeed, examination of transcriptomic profiles in white prepupae revealed a striking downregulation of ribosomal genes in *JHEH TKO* in both sexes but especially females (Fig 6c). Given well-established roles for ribosomal RNAs in both growth and developmental timing (KONGSUWAN *et al*. 1985) (LAMBERTSSON 1998), these data suggest that the requirement for JHEHs in growth is through impacts on ribosomal biology, while the requirement for JHEHs in timing is through regulation of the JH/ecdysone axes.

**Figure 6:**
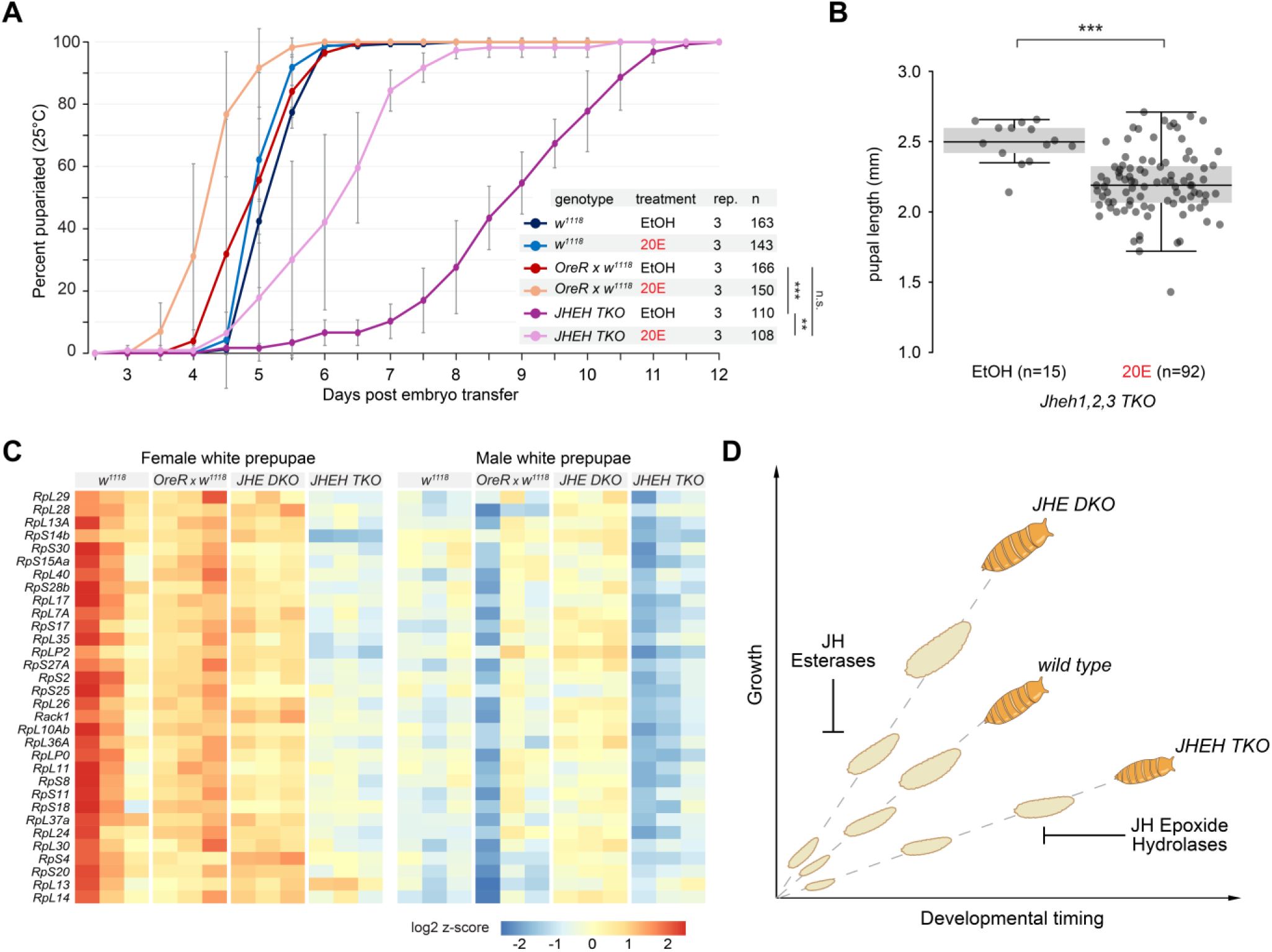
Defects in developmental timing, but not growth, in the absence of JHEHs is linked to dysregulated ecdysone signaling. **A** Graph of the percent pupariation relative to days post egg lay in control and *JHEH TKO* fed either the active form of ecdysone (20- Hydroxyecdysone or 20E) dissolved in ethanol (EtOH) or fed an equivalent amount of vehicle EtOH. Each dot indicates mean percent pupariated and whiskers represent standard deviation. The number of biological replicates (rep) and total sample size (n) are shown to the right of each genotype. All 50% pupariation data was normally distributed and a two-tailed Student’s t-test was used to test for statistical significance (* = p<0.05, ** = p<0.01, *** = p<0.001, n.s. = not significant). **B** Graph of pupal length in *JHEH TKO* animals after feeding either E20 or ethanol (EtOH) as a control 72 hours after egg lay. Dots indicate individual pupa, and sample size for each group is shown under the respective box plot. Statistical significance was tested by two- tailed Student’s t-test. P-value *** = <0.001. **C** Heatmap of differentially expressed ribosomal RNAs in white prepupae detected by RNA-sequencing. Genes are noted to the left, genotype and sex (M or F) is noted above each column, and heat map scale is below. **D** Schematic summarizing the working model based on data from this study, in which JHE restrain growth during development, while JHEHs restrict the time it takes to undergo developmental transitions from larvae (light beige) to pupae (brown).

## Discussion

Here, we investigated programmed hormone degradation using the JH degradation enzymes, JHEs and JHEHs, in *Drosophila melanogaster* as a model. *In vivo* requirements for these enzymes in this classic model organism are poorly understood due to paralogue expansion. By generating animals lacking both JHEs or all three JHEHs, we uncovered a separation of requirements for each programmed degradation pathway in two classic JH functions: developmental growth and timing. Despite distinct developmental requirements, we found that both types of JH catabolic enzymes restrain JH signaling as well as other metabolic gene networks and regulate endocrine gland morphology. Through our comparative approach, we found that JHEHs, but not JHEs, are integrated into several homeostatic feedback mechanisms, which together with more profound developmental requirements, suggest JHEHs are more critical to JH regulation during *D. melanogaster* juvenile development. As described further below, these results provided new insights into the complexity of programmed JH degradation and more broadly uncovered new aspects of JH biology and homeostasis.

In many insects, juvenile growth is linked to the timing of developmental transitions, with ectopic JH associated with increased body size and delayed metamorphosis. Whether *D. melanogaster* is susceptible to these classic JH effects has been less clear due to extensive genetic redundancies in JH synthesis enzymes and receptors (BURTENSHAW *et al*. 2008; ABDOU *et al*. 2011; JINDRA *et al*. 2015; FUJINAGA *et al*. 2026). Our study suggests that excess endogenous JH impacts this classic model organism like other insects, however growth and developmental timing phenotypes were genetically separable. Loss of JHEs did not delay development but had a mild impact on body size consistent with ectopic JH (MIRTH *et al*. 2014). In contrast, loss of JHEHs caused a severe developmental delay and reduced body size (Fig 3). Our findings are consistent with emerging patterns from non-Drosophila studies in which JHEs are more commonly linked to growth and JHEHs are linked to developmental timing in several non- *Drosophila* insects (TUSUN *et al*. 2017; VASQUEZ *et al*. 2023; LEI *et al*. 2025; QU *et al*. 2026). Yet, the mechanisms underlying this uncoupling of classically JH-linked functions is unknown. Each enzyme may act on additional substrates, or the products of JH catabolism may have their own biological functions, both of which have been shown for JHEs and their JH acid products (ISMAIL *et al*. 1998; STEINER *et al*. 2017; HOPKINS *et al*. 2019). Alternatively, each pathway may regulate JH titers at different developmental stages. We favor the latter, as *Jhehs* are more broadly expressed than *Jhes* during juvenile development in *D. melanogaster* (Supp Fig 1) and the classic JH target, *Kr-h1*, was more compromised in *JHEH TKO* animals throughout the juvenile period (Fig 2). However, additional biochemical and tissue-specific functional genetic studies will be required to properly distinguish between these possibilities.

Our comparative transcriptomic analysis provided important clues about shared and unique developmental requirements for JHEs and JHEHs. *JHE DKO* white prepupae exhibited a downregulation of spermatogenesis genes (Fig S4), perhaps consistent with recent reports of local JH signaling in the male reproductive tract (WILSON *et al*. 2003; LYU *et al*. 2023; BOX *et al*. 2024; KUROGI *et al*. 2024; RAMESH *et al*. 2025). Intriguingly, *JHEH TKO* white prepupae exhibited a downregulation of ribosomal genes (Fig 6), which may underlie our observation that these mutant animals tend to be smaller. Indeed, mutants in ribosomal genes cause both a developmental delay and smaller body size (KONGSUWAN *et al*. 1985; MARYGOLD *et al*. 2007). We speculate that JHEHs indirectly regulate ribosomal genes through JH titers as JH signaling has previously implicated in ribosomal gene expression in the mosquito (WANG *et al*. 2017). Similarly, increases in genes encoding Cyp450s and GSTs found upon the loss of either JHEs or JHEHs (Fig 4), may reflect responses to elevated JH titers. Both Cyp450s and GSTs are involved in both hormone synthesis as well as metabolism and detoxification of many small molecules (ENAYATI *et al*. 2005; ENYA *et al*. 2014). Application of ectopic JH or JH agonists induce Cyp450 and GSTs expression (ZHOU *et al*. 2006; WU AND LU 2008; DAVID *et al*. 2013; TARHAN *et al*. 2013), suggesting these genes may be a shared signature of elevated JHs. Induction of these genes may also reflect the existence of alternative JH degradation mechanisms. Indeed, Cyp4C7 was shown to convert JH III to 12-trans-hydroxy JH III and expression of this Cyp450 was induced after bursts of JH synthesis (SUTHERLAND *et al*. 1998).

Collectively, these findings suggest that investigations shed light on diverse biological responses to excess hormones.

This investigation into JH degradation pathways also uncovered several new layers of feedback within the JH axis. Loss of JHEH-mediated degradation led to not only upregulation of RNAs from the *Jheh3* promotor, but also five ecdysone synthesis genes (Fig 5). Interestingly, while ecdysone synthesis genes are only upregulated in *JHEH TKO* animals, both *JHE DKO* and *JHEH TKO* animals had enlarged prothoracic glands (Fig 5). These data are consistent with previous reports of the JH-responsiveness of the prothoracic gland and the broader ring gland (LIU *et al*. 2018), though results presented here demonstrate that loss of JH degradation increases, rather than decreases ring gland size. Whether these opposing outcomes are due to differences in genetic perturbations predicted to increase JH signaling, food composition, or another parameter is unknown. Interestingly, we did not find evidence of feedback between JHE and JHEH pathways, as has been shown in other insects (ZHANG *et al*. 2017; GLASTAD *et al*. 2020; LI *et al*. 2022), which may reflect this study’s transcriptomic dataset focuses on a narrow, 15-minute-long developmental stage. Future studies will be required to determine whether cross-regulation exists between the two pathways in other life stages or biological contexts in *D. melanogaster*. Nonetheless, our findings add to a growing appreciation for complexity homeostatic feedback within the JH axis (ABDOU *et al*. 2011; LEBOEUF *et al*. 2018; LEYRIA *et al*. 2022a; LEYRIA *et al*. 2022b; SMYKAL AND DOLEZEL 2023; BARTON *et al*. 2024).

Of the two JH degradation pathways, JHEH loss resulted in both a more severe developmental phenotype than JHE loss and robust activation of feedback. Together, these data suggest that JHEH-mediated JH degradation is the dominant pathway restraining JH titers during *D. melanogaster* juvenile development. This conclusion is significant in light of foundational studies in other insects that suggest circulating JHEs are thought to be the principal JH degradation pathway (KHLEBODAROVA *et al*. 1996). Indeed, increased JHE expression upon ectopic JH was historically used as a sensitive readout for JH titers (VENKATESH 1990). However, our findings are consistent with earlier studies showing that JHEH activity is much higher in *D. melanogaster* just prior to metamorphosis (KHLEBODAROVA *et al*. 1996). Consistent with JHEH pathway dominance, our data also suggest *bona fide* genetic redundancies among *Jhehs* in *D. melanogaster*. Indeed, a recent study found that loss of *Jheh1* and *Jheh2* caused only a four-hour delay in pupariation (ROGALSKI *et al*. 2025), while we find that loss of all three Jhehs caused a 5–7-day delay (Fig 3 and 6). While *Jheh3* may encode the ancient gene, all *D. melanogaster* Jhehs possess the required amino acids to catabolize JH and share similar expression profiles during development (Supp Fig 1). Whether the JHEH pathway dominance that we uncovered is restricted to juvenile growth and developmental timing or underappreciated genetic redundancies in *Jhehs* in other species will be an important area of future exploration.

Here, we defined the requirements for JH degradation enzymes in classic JH-functions: developmental timing and growth. However, JHs also have essential roles in reproduction, diapause, behavior, aging, as well as the establishment of sexual dimorphisms (GOTOH *et al*. 2011; YAMAMOTO *et al*. 2013; WU *et al*. 2018; SANTOS *et al*. 2019; EASWARAN *et al*. 2022).

Interestingly, transcriptomic data from this study suggest females may require programmed JH degradation more than males (Fig 4 and 6). Moreover, JH degradation enzymes are both responsive to insecticides and may be key regulators of insecticide resistance (HUANG *et al*. 2020). The mechanisms by which JH degradation controls various functions beyond juvenile growth and developmental timing are unclear; this study and the genetic tools generated in the tractable *Drosophila melanogaster* model provide a platform to shed light on these mechanisms. More broadly, results from this study demonstrate that such degradation enzymes can serve as molecular handles to uncover new aspects of hormone biology.

## Author Contributions

RS: Conceptualization, Investigation, Data analysis, Supervision, Writing - review & editing; KG: Methodology, Reagent generation, Investigation, Data analysis and Validation, Writing – review & editing; HS: Methodology, Reagent generation, Investigation, Data analysis; CH: Investigation, Data Analysis; IA: Investigation; AJ: Investigation, Data analyses; YN: Investigation; JY: Investigation; AAS: Data analysis, Supervision; LJB: Conceptualization, Formal analysis, Funding acquisition, Investigation, Project administration, Supervision, Validation, Writing - Original draft preparation, Writing - review & editing

## Data and Resource Availability

All relevant data and resources generated from this study can be found within the article and its supplemental material. RNA-Seq data have been uploaded to the Gene Expression Omnibus database (Accession Number GSE286242).

## Acknowledgments

We thank Dr. Brian Brigham for statistics consultation and members of the Barton Lab and Dr. Pamela Geyer for feedback. We thank Dr. Ruth Lehmann for support during the beginning of this project. Stocks obtained from the Bloomington Drosophil*a* Stock Center (NIH P40OD018537) were used in this study.

## Study funding

This work was supported by R00 HD097306 from the Eunice Kennedy Shriver National Institute for Child Health and Development to LJB. This research was conducted while Lacy J Barton was an AFAR Grant for Junior Faculty awardee. Lacy Barton, PhD holds a Voelcker Fund Young Investigator Award from the MAX AND MINNIE TOMERLIN VOELCKER FUND.

## Conflict of interest

Authors have no conflicts of interest to report.

---

**Supplemental Figure 1:**
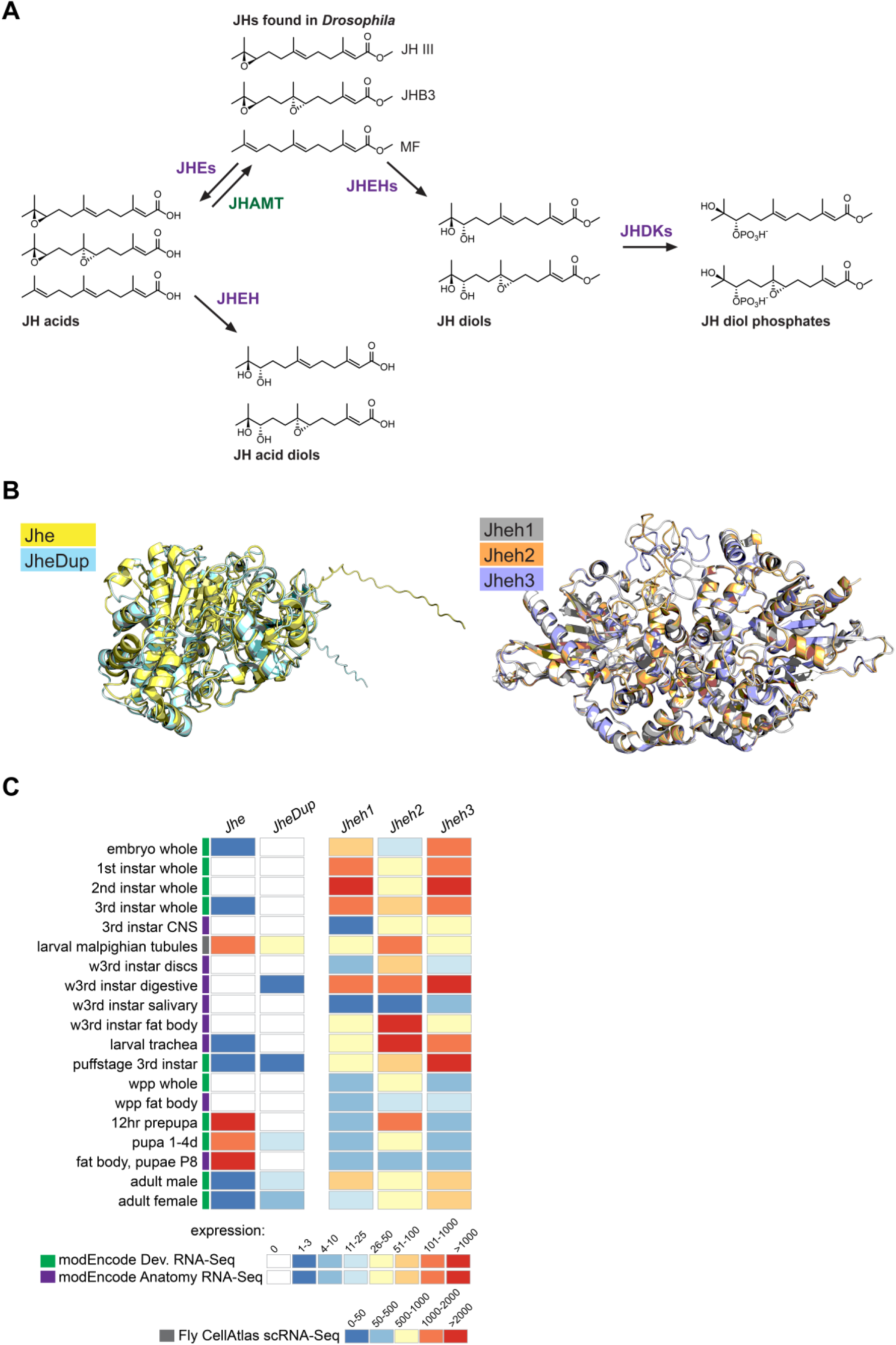
Data related to Figure 1: **A** Chemical structures of JHs found in *D. melanogaster*, as well as the products of JHE and JHEH catabolism. Also shown is conversion of JH acids to JHs by JHAMT. **B** Structural similarity of predicted structures for JH degradation enzymes. *Top*: Jhe (yellow) and JheDup (cyan), *Bottom*: Jheh1,2,3 (grey, orange, blue respectively). Both enzyme classes show significant structural overlap between paralogs. **C** Expression of JH degradation enzymes across *Drosophila* development and tissues. w3rd – wandering third instar larvae, wpp – white prepupal stage.

**Supplemental Figure 2:**
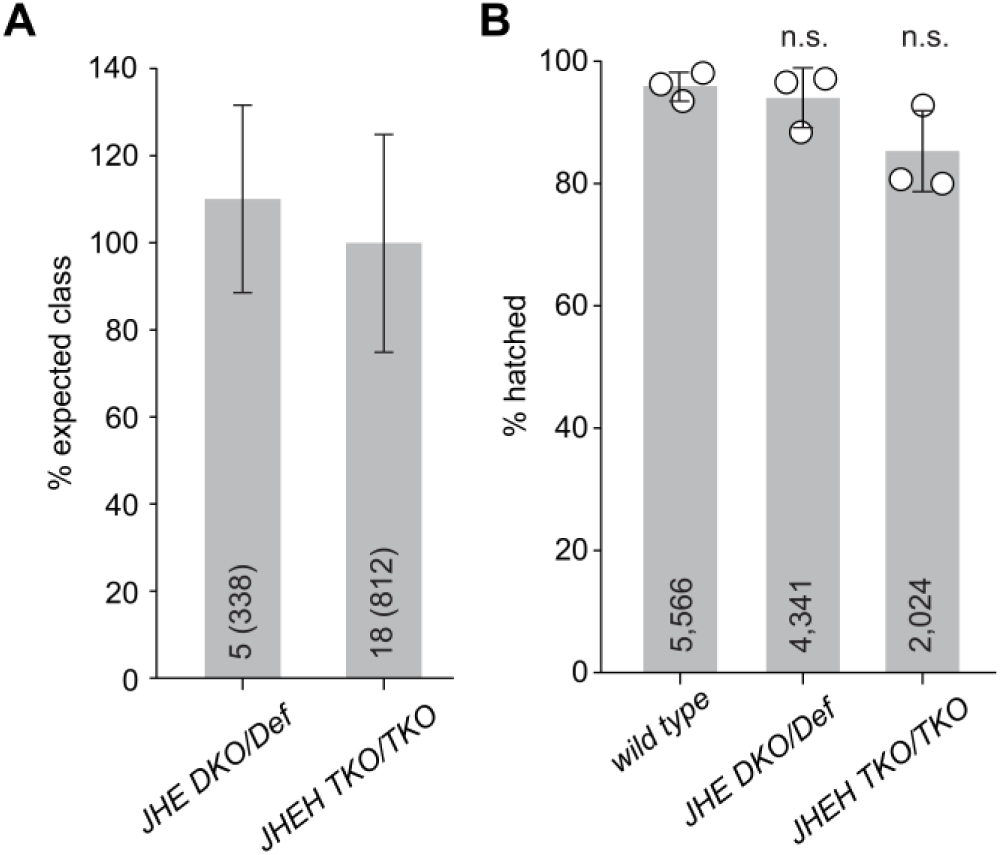
Data related to Figure 2: **A** Percent expected class for *JHE DKO* and *JHEH TKO* animals. The number of replicates is shown at the bottom of each bar, with total animals quantified in parentheses. Bars and whiskers represent mean ± SD. **B** Graph of hatch rate for JH degradation enzyme mutants. The maternal (M) and zygotic (Z) genotype for each bar is at the bottom of the graph. Each dot represents a biological replicate, and the total number of embryos analyzed is shown at the base of each bar. Bars and whiskers represent mean ± SD and significance tested by two-tailed Student’s *t*-test.

**Supplemental Figure 3:**
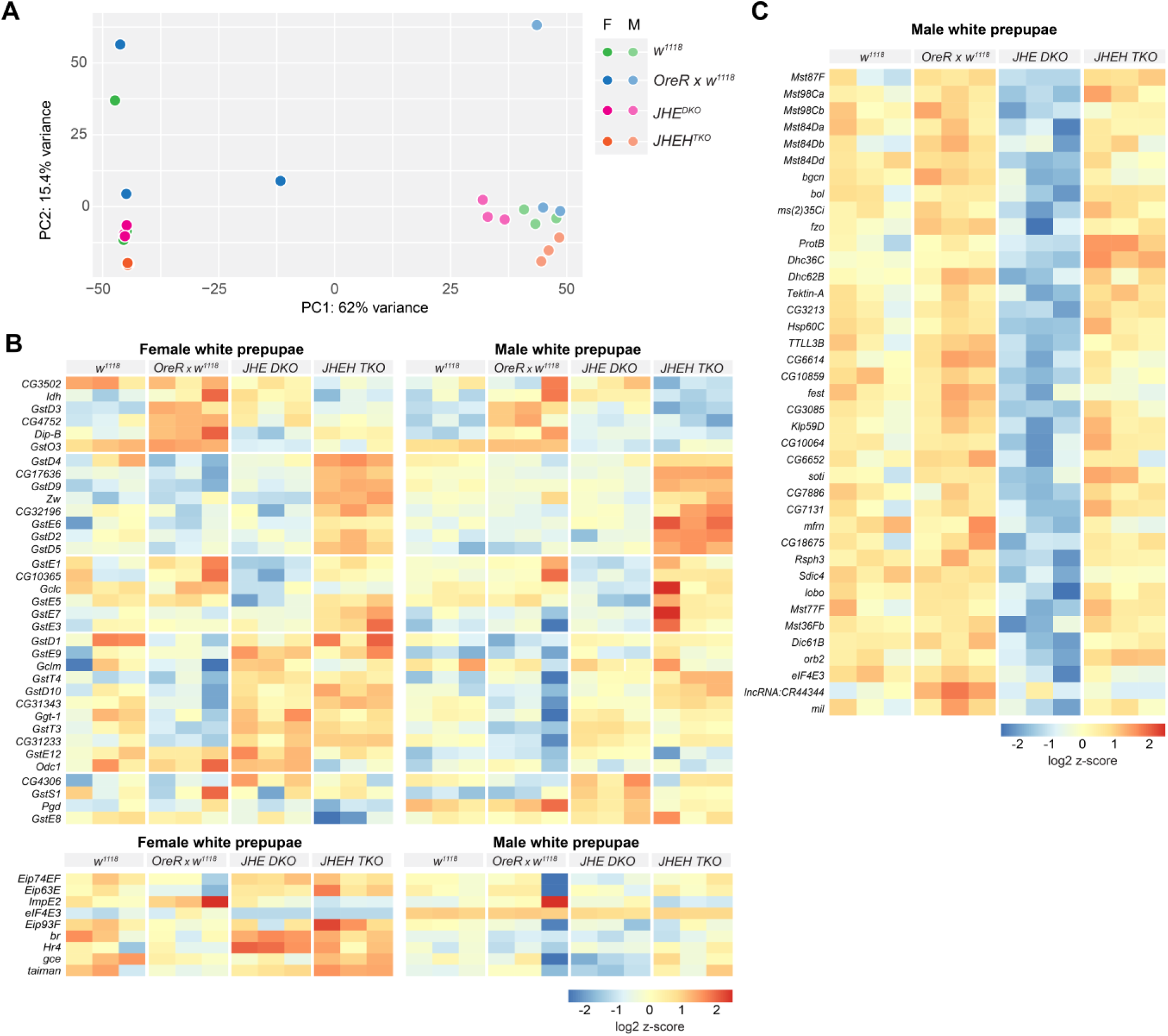
Data related to Figure 4: **A** Principal component analysis of bulk RNAseq of three biological replicates of whole white prepupae. **B** Heatmap of differentially expressed glutathione s-transferase (GST) and associated glutathione metabolism genes (top), ecdysone and JH-regulated genes (bottom). Genes are noted to the left, genotype and sex (M or F) is noted above each column, and heat map scale is below. **C** Heatmap of spermatogenesis genes (bottom) in male white prepupae, with scale is below.

**Supplemental Figure 4:**
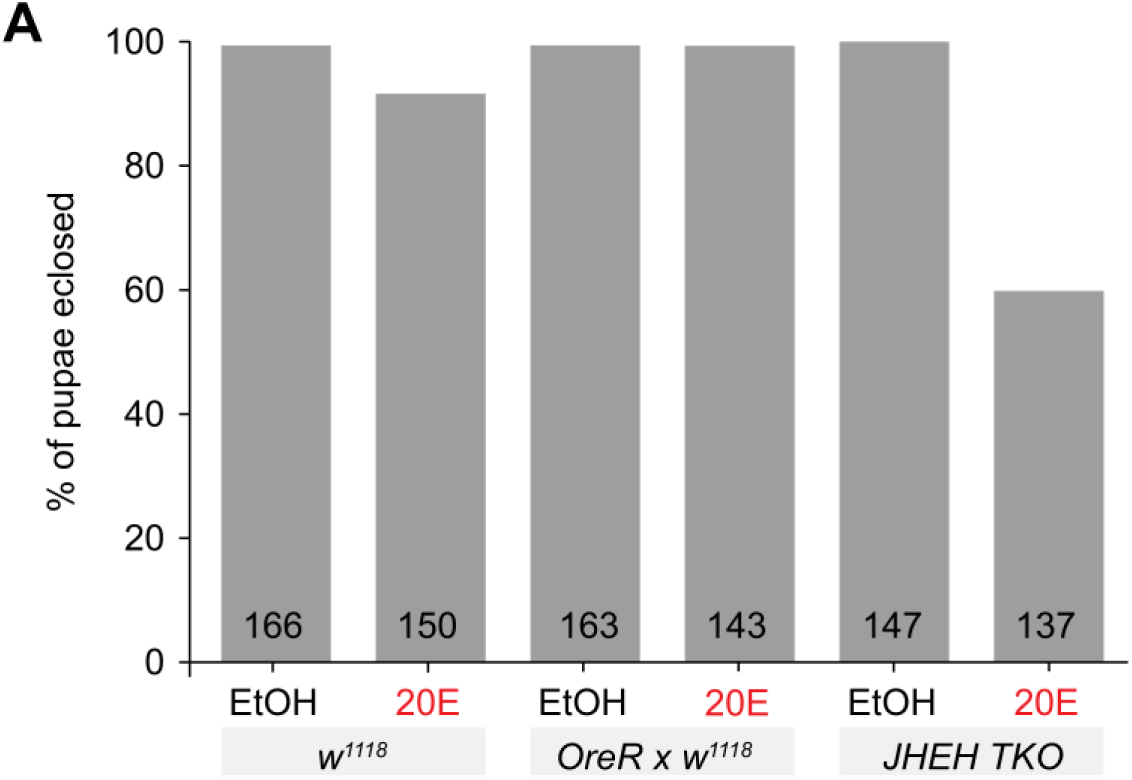
Data related to Figure 5: Graph of percent eclosion after either feeding of food containing ethanol solvent control (EtOH) or 20-Hydroxyecdysone (20E, red) 72 hours after egg lay as described in the Methods. Genotypes and treatment represented by each bar is shown at the bottom of the graph. The number of animals quantified is shown at the base of each bar.

**Supplemental Table 1.**
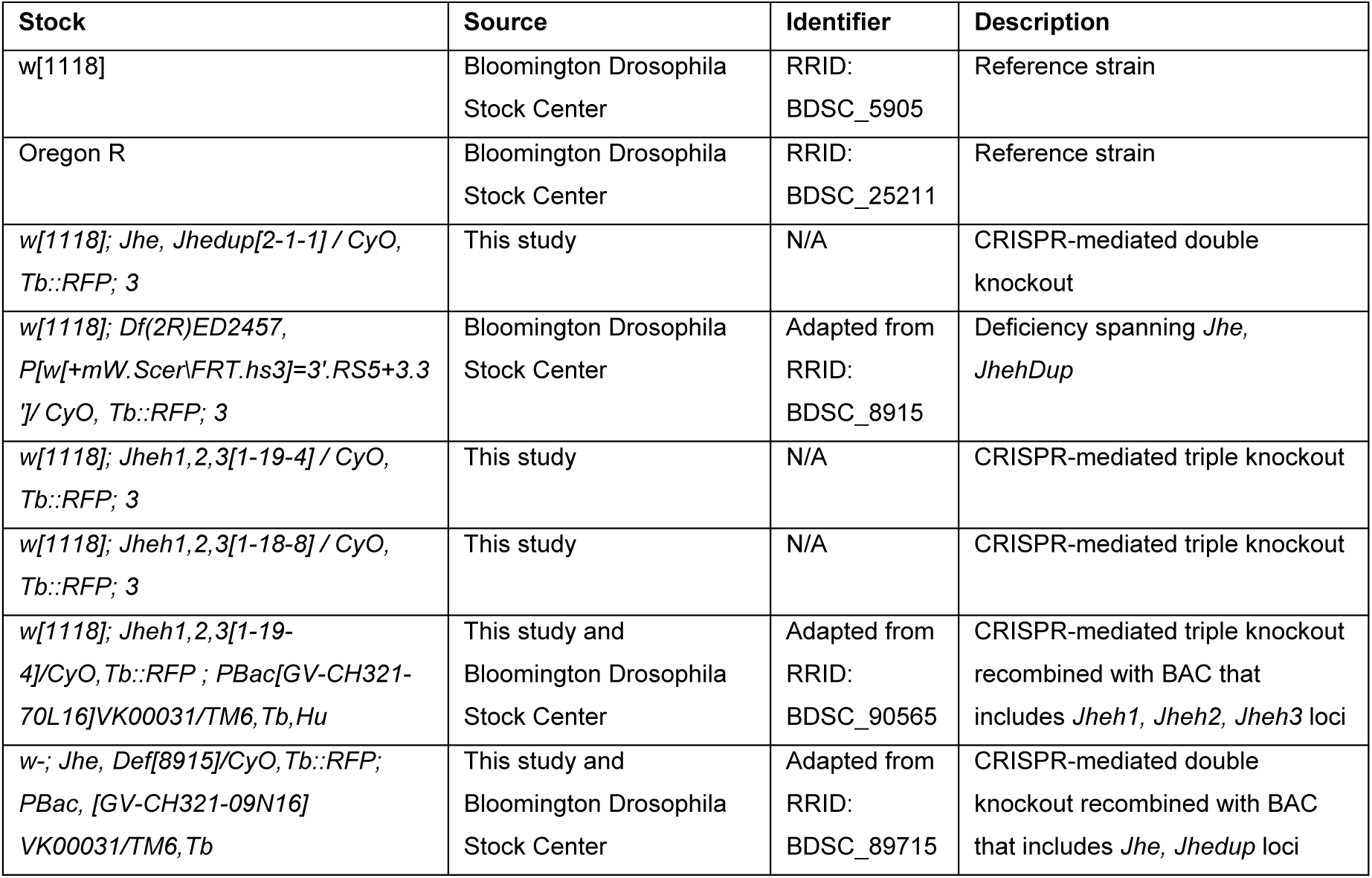
*Drosophila* strains used in this study.

**Supplemental Table 2.**
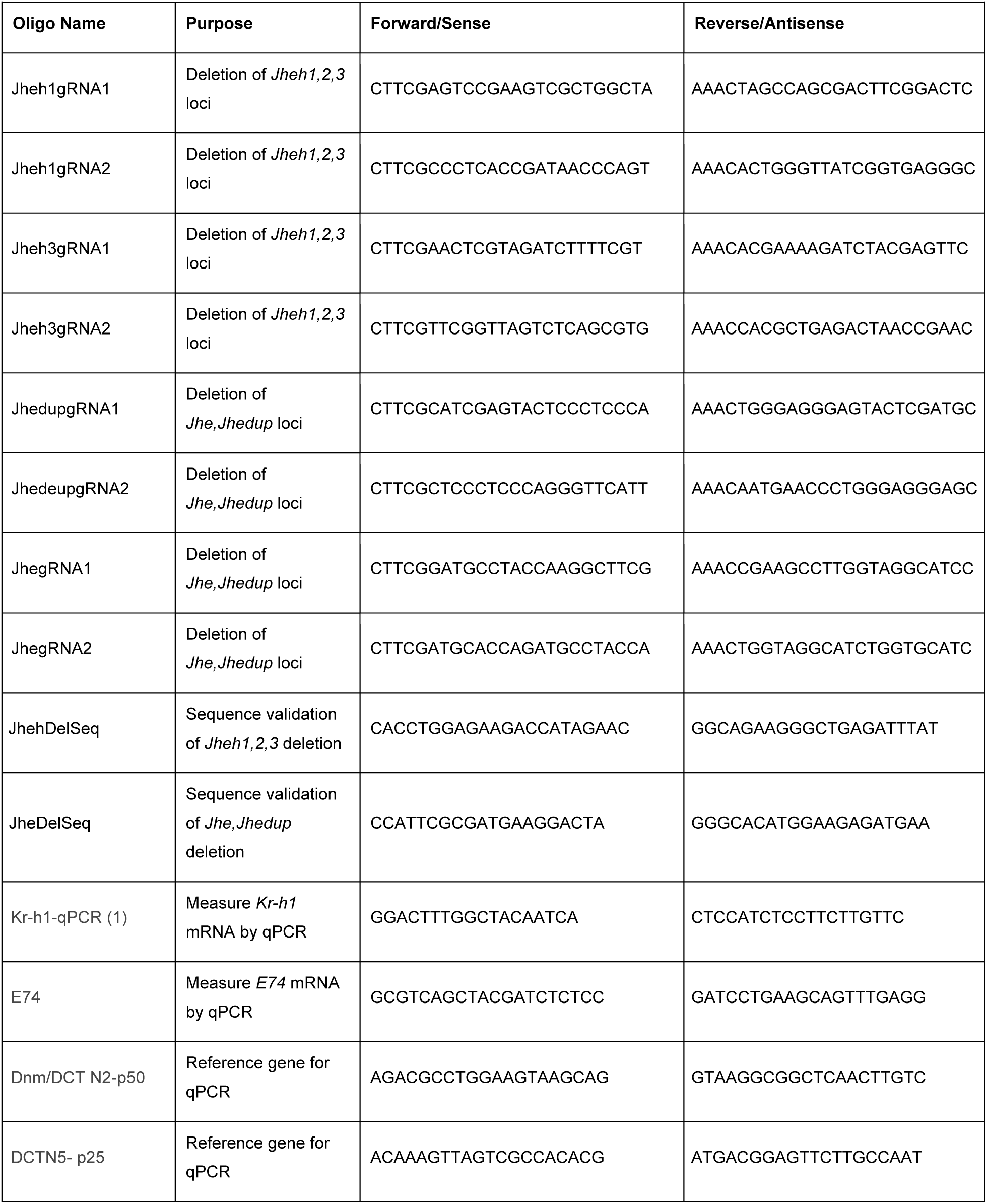
Oligonucleotides used in this study.

## Supplemental Files

**DatasetS1_MaleDEGsWPP:** Differentially expressed genes in *JHE DKO* and *JHEH TKO* male white prepupae.

**DatasetS2_FemaleDEGsWPP:** Differentially expressed genes in *JHE DKO* and *JHEH TKO* female white prepupae.

**DatasetS3_GOTerms:** Gene ontology analyses of differentially expressed genes in *JHE DKO* and *JHEH TKO* white prepupae.

## Notes

### Competing Interest Statement

The authors have declared no competing interest.

### Summary of Updates

Additional experiments and figures. Authors added.

